# Multi-hit STAG2 mutations define a high-risk subset of MDS and reveal convergent evolutionary targeting of cohesin

**DOI:** 10.64898/2026.09.02.749005

**Authors:** Eno-obong B. Udoh, Yi Chen, Matteo D’Addona, Edna Stewart, Rong Deng, Felipe de Almeida Sartori, Serhan Unlu, Zachary Brady, Ruowei Zhu, Xiaoyi Cheng, Viviana Scoca, Jane J. Xu, Sergei Doulatov, Karl Theil, David Bosler, Valeria Visconte, Jaroslaw P. Maciejewski, Aaron D. Viny

## Abstract

STAG2 is the most frequently mutated cohesin gene in myeloid neoplasms, yet the significance of multiple mutations within this X-linked tumor suppressor remains unknown.

We analyzed a cohort of 1,967 adult patients with myeloid neoplasms and identified 233 cases (12%) harboring STAG2 mutations, including 38 cases (16%) with multiple STAG2 hits. Patients with multi-hit STAG2 mutations exhibited increased multilineage dysplasia compared with single-hit cases and experienced inferior overall survival, an effect driven primarily by patients with myelodysplastic syndromes (MDS).

To investigate the molecular basis of recurrent STAG2 acquisition, we performed long-read sequencing in representative cases with phaseable STAG2 mutations. In the informative case examined, distinct truncating STAG2 mutations did not co-occur on the same DNA molecule, supporting independent acquisition rather than stepwise allelic inactivation. Cohort-level variant allele frequency patterns were consistent with recurrent evolutionary targeting of STAG2 across related clonal populations.

Together, these findings support a model in which multi-hit STAG2 mutations arise through convergent evolution and define a biologically distinct, adverse-risk subset of MDS.

**Key points:**

- Multi-hit *STAG2* mutations identify a high-risk subset of MDS characterized by increased multilineage dysplasia and inferior survival.
- Long-read sequencing supports convergent evolutionary acquisition of *STAG2* mutations rather than stepwise allelic inactivation.

## Introduction

Myelodysplastic syndromes (MDS) are clonal hematopoietic disorders characterized by ineffective hematopoiesis, morphologic dysplasia, and a variable risk of progression to acute myeloid leukemia (AML)^1–4^. Disease evolution is driven by the stepwise acquisition of somatic mutations that establish clonal dominance and shape clinical phenotypes. Among recurrently mutated genes, components of the cohesin complex are enriched in higher-risk MDS and secondary AML, implicating altered chromatin organization as a central feature of myeloid transformation ^5–9^.

Somatic mutations in the core cohesin complex genes, including *STAG2*, *RAD21*, *SMC1A*, and *SMC3,* are frequently observed in myeloid neoplasms (MNs)^4^. Among these, *STAG2* is the most commonly mutated and is predominantly affected by truncating, loss-of-function alterations, consistent with tumor suppressor activity ^10^. Located on the X chromosome, *STAG2* is typically considered a monoallelic tumor suppressor in hematopoietic cells, as a single inactivating mutation is sufficient to disrupt cohesin function in the hemizygous state^11^. Clinically, *STAG2* mutations are associated with adverse outcomes and frequently precede overt leukemia, suggesting a role in early clonal selection. In addition, mutations in *STAG2* often predict a previously undiagnosed antecedent MDS in patients newly diagnosed with AML^4,12^. Functionally, *STAG2* loss alters chromatin looping, transcriptional insulation, and lineage specification^7^.

Despite this framework, a subset of patients harbors multiple independent *STAG2* mutations. In contrast to autosomal tumor suppressors, such as *TP53* or *TET2*, in which multiple hits often reflect biallelic inactivation, the biological significance of multiple mutations in an X-linked gene remains unclear^13,14^. Specifically, it remains unclear whether recurrent STAG2 mutations arise through convergent evolution in independent subclones or through sequential acquisition within a single allele^11^. Given the essential role of cohesin in hematopoietic stem cell function, we hypothesized that multi-hit STAG2 states reflect selective pressure for dose-constrained cohesin dysfunction achieved through independent evolutionary acquisition of STAG2 mutations rather than stepwise allele-restricted evolution.

To test this, we analyzed a large cohort of patients with *STAG2*-mutant myeloid neoplasms, integrating clinical, genomic, and clonal hierarchy analyses. In representative cases, we applied long-read nanopore sequencing to directly determine the allelic configuration. Here, we demonstrate that multi-hit *STAG2* mutations define a clinically distinct subgroup enriched for adverse features in MDS and provide evidence supporting independent acquisition of STAG2 mutations through convergent evolution. These findings establish allelic configuration as an important dimension of *STAG2* biology and suggest that recurrent perturbation of cohesin function contributes to disease severity^15^.

## Materials and methods

### Patients

This study was approved by the Cleveland Clinic and Columbia University Irving Medical Center Institutional Review Boards (IRB). A total of 1,967 adult patients within our myeloid neoplasm (MN) cohort at the Cleveland Clinic were screened for the presence or absence of *STAG2* mutations of any kind identified on next-generation sequencing (NGS) of periphera blood or bone marrow. A complete list of genes covered by the Cleveland Clinic hematologic neoplasm NGS panel is available in eTable 1. Patients were further classified based on the number of *STAG2* variants detected. Patients with 1 *STAG2* mutation were classified as monoSTAG2^MT^, while those with >1 *STAG2* mutation were grouped as multiSTAG2^MT^. Clinical information on patient age, sex, diagnosis, karyotype, peripheral blood, and bone marrow was obtained from electronic medical records. Diagnosis was based on the 2016 World Health Organization (WHO) criteria for myeloid neoplasms and acute leukemia^16^.

### Assessment of clonal hierarchy

Clonal hierarchy was approximated for the first *STAG2* hit in the multiSTAG2^MT^ cohorts using the Pyclone pipeline, which was initially developed to infer clonal population structure in cancer and has also been validated in myeloid neoplasms cohorts of patients with *TET2* and *SF3B1* mutations^17–19^. This variant allele frequency (VAF)-based analytic method was used to assign clonal hierarchy to the first *STAG2* hit relative to other co-mutations. The 1^st^ STAG2 hit was designated as either ancestral (dominant or codominant) or subclonal/secondary which are further defined as follows: ancestral/dominant (DOM) if it had the highest VAF which was at least 5% greater than the second highest VAF, as ancestral/codominant (COD) if the difference between the highest and second highest was less than 5% VAF (includes cases where *STAG2* and another mutation are “tied” for the lead), and as subclonal/secondary (SEC) if there exists at least one mutation whose VAF is at least 5% higher than the VAF of the 1^st^ *STAG2* hit. Representative cases of patients with the 1^st^ *STAG2* hit as dominant, co-dominant, and subclonal are shown in Figure 1. This approach assumes comparable copy number states across mutations and may not fully capture clonal architecture in all cases.

**Figure 1.**
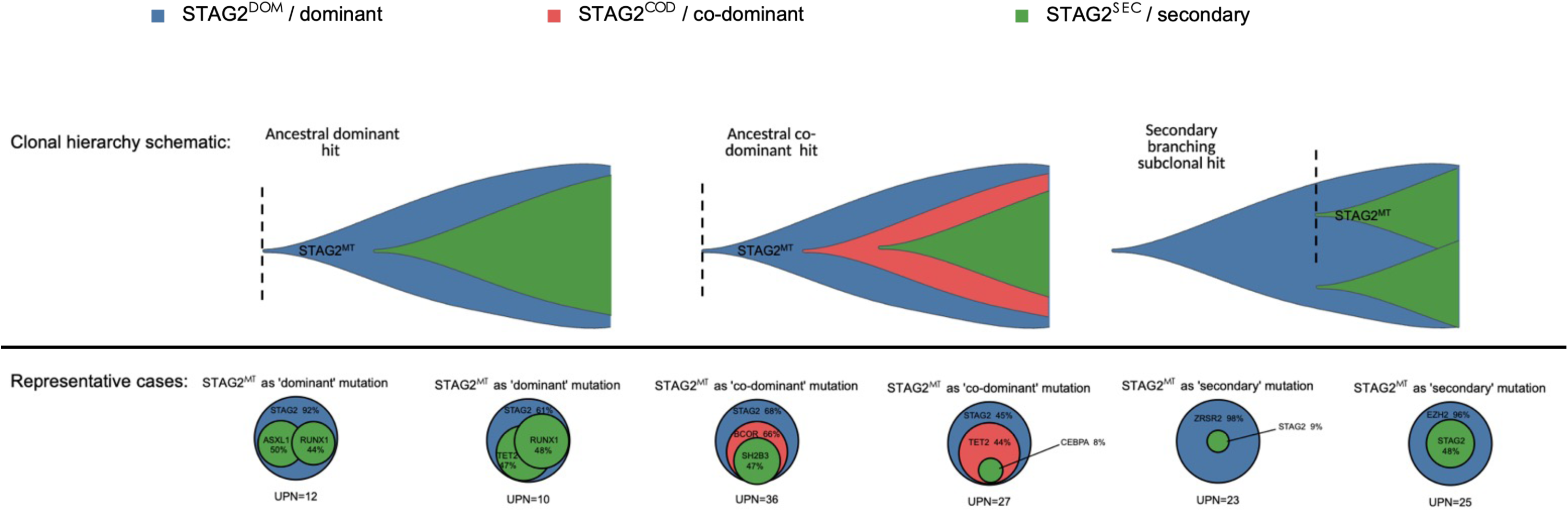
VAF-based approach for clonal hierarchy assessment of the 1^st^ STAG2 hit for multiSTAG2^MT^ cases. Top: Fish plot of conceptual patterns of STAG2 involvement during clonal evolution. Bottom: Pie chart of the representative patient cases shown for each clonal ranking highlighting the relative VAF relationships between the 1^st^ STAG2 hit and other co- occurring mutations.

### Assessment of hematologic parameters and bone marrow pathology

Hematologic parameters including white blood cells (WBC), hemoglobin (Hgb), platelet (Plt), and absolute neutrophil count (ANC) were further assessed for cytopenias and cytoses. Cytopenias were defined as follows: anemia (Hgb < 12g/dL for females, and Hgb < 13g/dL for males), neutropenia (ANC < 1.8K/uL) and thrombocytopenia (Plt < 150K/uL)^20^. Based on the number of cytopenias present, patients were then grouped as having “monocytopenia”, “bicytopenia” and “pancytopenia.” In contrast, cytosis was defined as follows: erythrocytosis (Hgb ≥ 17g/dL), neutrophilia (ANC ≥ 6.0 K/uL), and thrombocytosis (Ptl ≥ 450 K/uL)^21^. Bone marrow (BM) biopsies and aspirate smears were reviewed for blast percentage, cellularity, fibrosis, and evidence of dysplasia in the erythroid, myeloid, and megakaryocytic lineages. Patterns of dysplasia previously observed in STAG2^MT^ patients include but are not limited to abnormal myeloid nuclear segmentation and cell granulation (hypo- and hyper-granulation), separated megakaryocyte nuclear lobes, and erythroid nuclear irregularities including binucleation^22,23^. Select cases from our cohort were reviewed by an independent pathologist for dysplastic features. Patients were further graded by the number of lineages found to be dysplastic: “unilineage,” “bilineage,” and “trilineage”. Lastly, BM cellularity was defined as hypercellular (≥ 70%), normocellular (30-70%), and hypocellular (≤ 30%).

### Long-read sequencing for allelic phasing

For a subset of patients with available genomic DNA, long-read sequencing with adaptive sampling (PromethION, Oxford Nanopore Technologies) was performed to assess allelic configuration of multiple *STAG2* mutations. Samples were selected based on the genomic distance between mutations, prioritizing cases in which variants were separated by ≤10–20 kb to enable coverage within a single sequencing read. DNA quality and fragment size were assessed using Tapestation (Agilent) and Qubit (Thermo Fisher Scientific) before library preparation. Long-read sequencing enables direct observation of co-occurring variants within individual DNA molecules, allowing determination of whether mutations reside on the same allele (cis) or on different alleles or subclones (trans). Two patients (UPN32 and UPN33) met these criteria and were analyzed for read-level phasing.

### Statistical analysis

Continuous variables were compared using Student’s t-test and Wilcoxon rank-sum test, as appropriate. Categorical variables were compared using chi-square or Fisher’s exact tests. Overall survival was estimated using the Kaplan–Meier method and compared using the log-rank test. For analyses of co-mutation frequencies across genes, p-values were adjusted for multiple comparisons using the Benjamini-Hochberg false discovery rate (FDR) method. Two-sided p-values <0.05 were considered statistically significant. All statistical analyses were performed using SAS® and R®.

## Results

### Multihit STAG2 mutations are enriched for truncating variants without evidence of complete gene loss

Within a cohort of 1,967 patients with MN, we identified 233 cases (12%) harboring *STAG2* mutations. Of these, 84% (n = 195) of patients had 1 *STAG2* mutation (monoSTAG2^MT^), and 16% (n = 38) had more than 1 *STAG2* mutation (multiSTAG2^MT^). Within the multiSTAG2^MT^ cohort, 31/38 patients had 2-hits, 5/38 patients had 3-hits, and 2/38 had 4-hits (Figure 2A). All multiSTAG2^MT^ patients had at least one truncating mutation (nonsense, splice-site, or frameshift), with only 3 cases harboring both missense and truncating mutations. In contrast, monoSTAG2^MT^ cases were represented by 77% truncating mutations and 19% missense mutations (Figure 2B). The predominance of truncating alterations across multi-hit cases, without evidence for complete allelic deletion, is consistent with recurrent selection for partial STAG2 dysfunction without evidence for complete allelic loss, suggesting selection against full STAG2 inactivation.

**Figure 2.**
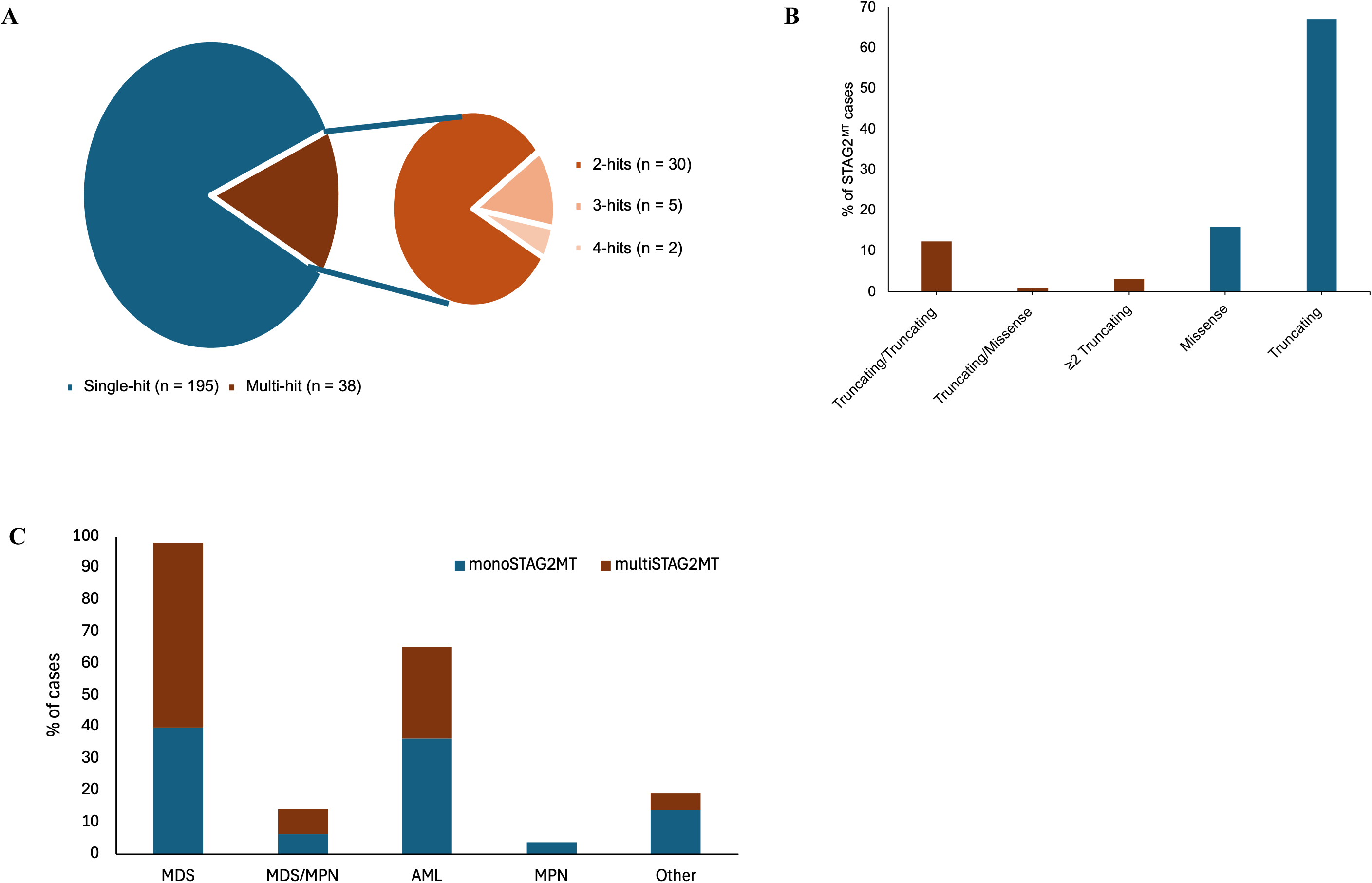
STAG2^MT^ classification and variant characteristics. **A)** Pie chart showing the breakdown of the number of mutations within the STAG2^MT^ cohort. **B)** Bar graph of the protein consequences (truncating: nonsense, frameshift, insertion/deletions, or missense) of STAG2 mutations present within the single-hit and the multi-hit groups. **C)** Bar graph showcasing the distribution of single-hit and multi-hit STAG2 cases across different MN subtypes.

*multiSTAG2^MT^ cases harbor a greater degree of dysplasia compared to monoSTAG2^MT^ cases* Assessment of clinical phenotype revealed that multiSTAG2^MT^ cases were diagnosed at a similar age to monoSTAG2^MT^ (p = 0.67) (eTable 2). 79% of patients in the multiSTAG2^MT^ cohort were male compared to 64% in monoSTAG2^MT^ (p = 0.07). Based on the 2016 WHO classification, 58% of multiSTAG2^MT^ patients were diagnosed with MDS, 8% with MDS/MPN, 29% with AML (21% s-AML, 5% pAML, and 3% t-AML), and 5% with CH/CCUS. Within the monoSTAG2^MT^ group, 40% of patients were diagnosed with MDS, 6% with MDS/MPN, and 36% with AML (18% s-AML, 17% pAML), and 14% with CH/CCUS (Figure 2C). There was no difference in the distribution of diagnosis across both cohorts (p = 0.30). The proportion of patients harboring normal versus abnormal cytogenetics was comparable across both cohorts (p = 0.50). In addition, there was no difference in the hematologic parameters for both cohorts, including assessment of cytopenia (eTable 3). Bone marrow morphology assessment showed that there was a greater degree of dysplasia in the myeloid (p = 0.04) and megakaryocytic lineages (p = 0.02) for patients with the multiSTAG2^MT^ cohort and a greater degree of dysplasia in more than one lineage (p = 0.02) compared to the monoSTAG2^MT^ group. These findings are consistent with impaired lineage fidelity associated with increased cohesin perturbation.

### Multi-hit STAG2 mutations are associated with higher-risk MDS features but not AML

Analysis of STAG2^MT^ patients by disease subtype: AML vs. MDS showed that there was no difference in the demographic features (age, sex) for these patients regardless of whether they harbored single or multiple hits in *STAG2* (p > 0.05); the male: female (M: F) ratio was 3.4 in MDS patients with multiSTAG2^MT^ versus 4.5 in AML patients with multiSTAG2^MT^. However, there was a significant difference in the MDS subtypes between both *STAG2* groups (Table 1). MDS patients with multiple hits were likely to have the MDS-EB-2 subtype compared to MDS patients with single *STAG2* hits (45.5% vs. 16.7%, p = 0.0046). This correlated with a greater blast % within the multi-hit *STAG2* MDS cohort compared to those with single-hits (p = 0.04) (Table 2). In contrast, multiSTAG2^MT^ patients with AML were more likely to be diagnosed with secondary AML (sAML) compared to the monoSTAG2^MT^ AML group (72.7% vs. 50.7%, p = 0.03) (Table 3). Conversely, this group was less likely to be diagnosed with primary AML (pAML) compared to the monoSTAG2^MT^ group (18.2% vs. 32.4%, p = 0.03). Furthermore, blast % and myeloid/erythroid (M/E) ratio were also higher in the monoSTAG2^MT^ group with AML compared to the multiSTAG2^MT^ group (p < 0.05) (Table 4).

**Table 1.** Comparison of baseline characteristics in multiSTAG2^MT^ and monoSTAG2^MT^ MDS cases.

| **Variables** | **monoSTAG2<sup>MT</sup> (n = 78)** | **multiSTAG2<sup>MT</sup> (n = 22)** | **p-value** |
| --- | --- | --- | --- |
| **Age (years)** | 73.8 ± 9.0 | 70.8 ± 10.8 | 0.20 |
| ≥60 | 74 (94.9) | 20 (90.9) | 0.61 |
| <60 | 4 (5.1) | 2 (9.1) |  |
| **Sex** |  |  |  |
| Male | 49 (62.8) | 17 (77.3) | 0.20 |
| Female | 29 (37.2) | 5 (22.7) |  |
| M:F ratio | 1.69 | 3.40 |  |
| **WHO Classification (2016)** |  |  |  |
| **MDS** |  |  | **0.05** |
| MDS-EB-1 | 25 (32.1) | 5 (22.7) |  |
| MDS-EB-2 | 13 (16.7) | 10 (45.5) | **0.0046\*** |
| MDS-MLD | 28 (35.9) | 3 (13.6) | 0.0462 |
| MDS-MLD-RS | 4 (5.1) | 3 (13.6) |  |
| MDS-SLD | 4 (5.1) | 0 (0.0) |  |
| MDS-SLD-RS | 1 (1.3) | 0 (0.0) |  |
| MDS with isolated del(5q) | 1 (1.3) | 0 (0.0) |  |
| MDS-U | 2 (2.6) | 1 (4.6) |  |
| Progressed to AML | 14 (29.2) | 9 (40.9) | 0.33 |
| **Cytogenetics** |  |  |  |
| Normal karyotype | 40 (58.0) | 10 (52.6) | 0.68 |
| Abnormal karyotype | 29 (42.0) | 9 (47.4) |  |

Abbreviations: EB – excess blast, MLD – multilineage dysplasia, RS – ringed sideroblast, SLD – single lineage dysplasia, U – unclassified. \* - Bonferroni's correction for multiple comparisons.

**Table 2.** Comparison of clinical characteristics in multiSTAG2^MT^ and monoSTAG2^MT^ MDS cases.

| **Variables** | **monoSTAG2<sup>MT</sup> (n = 78)** | **multiSTAG2<sup>MT</sup> (n = 22)** | **p-value** |
| --- | --- | --- | --- |
| **Hematologic parameters** |  |  |  |
| WBC (10<sup>9</sup>/L) (median/range) | 2.76 (1.50, 4.20) | 2.52 (1.40, 5.67) | 0.93 |
| <4.5 x 10<sup>9</sup>/L - leukopenia | 59 (76.6) | 14 (63.6) | 0.22 |
| >11.0 x 10<sup>9</sup>/L - leukocytosis | 3 (3.9) | 1 (4.6) | 1.00 |
| Hemoglobin (g/dL) | 8.3 (7.6, 9.5) | 8.0 (7.5, 8.9) | 0.57 |
| <12 g/dL (F) or <13g/dL (M) - anemia | 77 (100) | 22 (100.0) |  |
| >17 g/dL - erythrocytosis | 0 (0.0) | 0 (0.0) |  |
| Platelet (10<sup>9</sup>/L) | 68 (39, 146) | 80 (39, 167) | 0.56 |
| <150 X 10<sup>9</sup>/L - thrombocytopenia | 58 (75.3) | 16 (72.7) | 0.80 |
| >450 X 10<sup>9</sup>/L - thrombocytosis | 1 (1.3) | 2 (9.1) | 0.12 |
| ANC (10<sup>9</sup>/L) | 1.09 (0.38, 2.49) | 0.96 (0.34, 2.67) | 0.84 |
| <1.8 x 10<sup>9</sup>/L - neutropenia | 55 (71.4) | 11 (57.9) | 0.25 |
| >6.0 x 10<sup>9</sup>/L - neutrophilia | 5 (6.5) | 0 (0.0) | 0.58 |
| **Cytopenias** |  |  |  |
| Monocytopenia | 6 (7.8) | 4 (18.2) | 0.36 |
| Bicytopenia | 25 (32.5) | 6 (19.4) |  |
| Pancytopenia | 46 (59.7) | 12 (54.6) |  |
| **Bone Marrow Morphology** |  |  |  |
| Blasts | 4.0 (2.0, 8.0) | 8.0 (3.0, 11.0) | **0.04** |
| ≥ 5% | 36 (46.2) | 14 (63.6) | 0.15 |
| Cellularity |  |  |  |
| Hypercellular | 36 (52.2) | 14 (70.0) | 0.08 |
| Normocellular | 19 (27.5) | 6 (30.0) |  |
| Hypocellular | 14 (20.3) | 0 (0.0) |  |
| M/E ratio | 2.00 (0.90, 4.30) | 1.84 (1.00, 3.50) | 0.97 |
| Myelofibrosis | 15 (20.8) | 3 (15.0) | 0.75 |
| **Dysplastic Lineage** |  |  |  |
| Myeloid | 41 (56.2) | 15 (75.0) | 0.13 |
| Erythroid | 58 (79.5) | 15 (71.4) | 0.55 |
| Megakaryocytic | 64 (87.7) | 20 (95.2) | 0.45 |
| **Number of dysplastic BM lineage** |  |  |  |
| None | 1 (1.4) | 0 (0.0) | 0.71 |
| Unilineage | 7 (9.6) | 2 (9.5) |  |
| Bilineage | 39 (53.4) | 9 (42.9) |  |
| Trilineage | 26 (35.6) | 10 (47.6) | - |

Abbreviations: F – female, M – male, M/E – myeloid/erythroid.

**Table 3.** Comparison of baseline characteristics in multiSTAG2^MT^ and monoSTAG2^MT^.

| **Variables** | **monoSTAG2<sup>MT</sup> (n = 71)** | **multiSTAG2<sup>MT</sup> (n = 11)** | **p-value** |
| --- | --- | --- | --- |
| **Age (years)** | 70.8 ± 13.7 | 73.6 ± 9.7 | 0.51 |
| ≥60 | 55 (77.5) | 10 (90.9) | 0.44 |
| <60 | 16 (22.5) | 1 (9.1) |  |
| **Gender** |  |  | 0.49 |
| Male | 48 (67.6) | 9 (81.8) |  |
| Female | 23 (32.4) | 2 (18.2) |  |
| M:F ratio | 2.09 | 4.50 |  |
| **WHO Classification (2016)** |  |  |  |
| **AML** |  |  | **0.03** |
| sAML | 35 (50.7) | 8 (72.7) |  |
| tAML | 0 (0.0) | 1 (9.1) |  |
| pAML | 34 (49.3) | 2 (18.2) |  |
| **Cytogenetics** |  |  |  |
| Normal karyotype | 35 (57.4) | 5 (45.4) | 0.52 |
| Abnormal karyotype | 26 (42.6) | 6 (54.6) |  |

Abbreviations: sAML – secondary AML, tAML – therapy-related AML, pAML – primary AML.

**Table 4.**
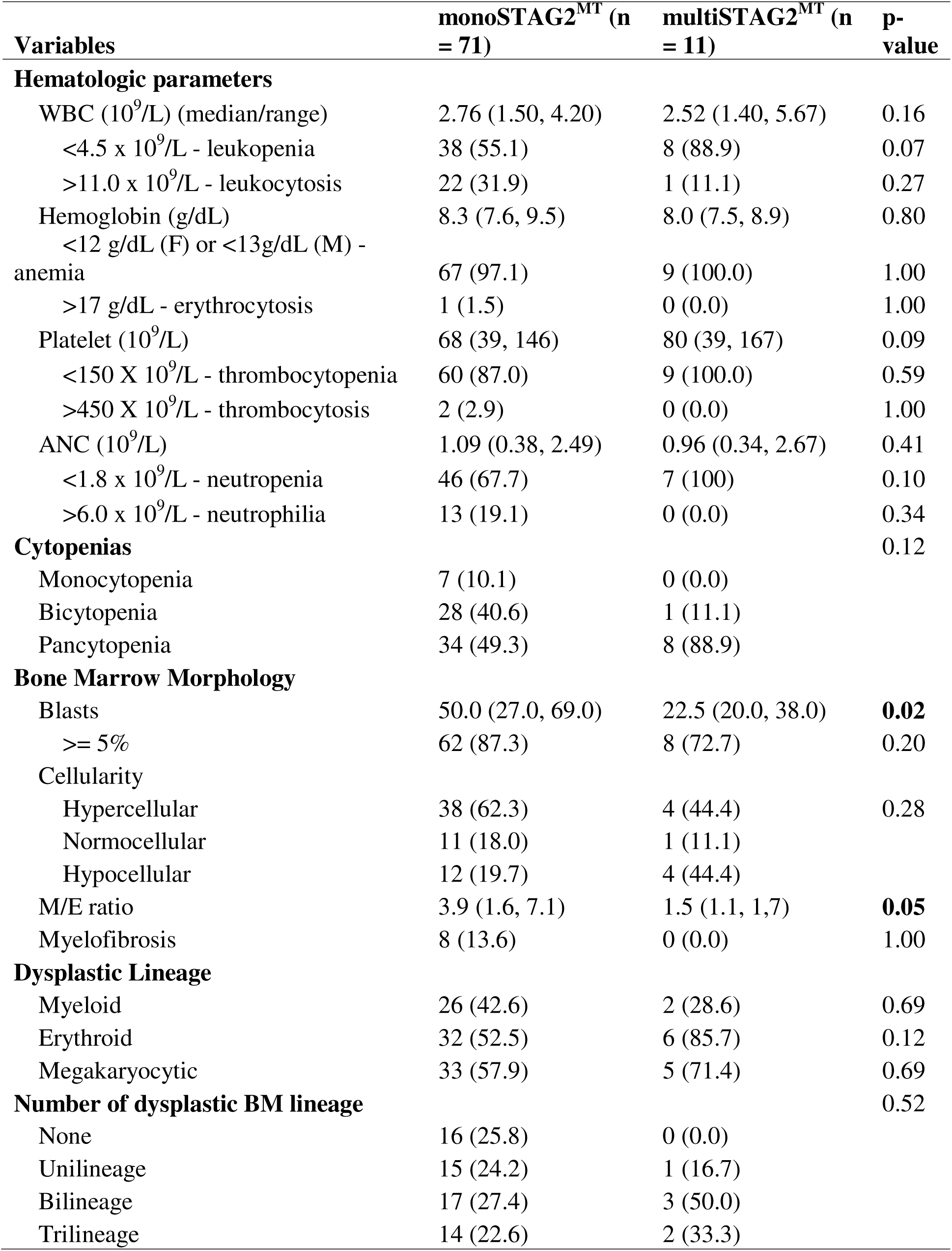
Comparison of clinical characteristics in multiSTAG2^MT^ and monoSTAG2^MT^.

### Distinct characteristics associated with multiSTAG2MT

To identify clinical features independently associated with multi-hit *STAG2* status, we performed univariable and multivariable analyses incorporating demographic, hematologic, cytogenetic, and morphologic variables (Figure 3). Across the entire cohort, patients with multiSTAG2MT exhibited increased dysplasia in the myeloid and megakaryocytic lineages, consistent with the increased burden of multilineage dysplasia observed in descriptive analyses. In contrast, age, peripheral blood counts, and cytogenetic status were not consistently associated with multi-hit status. When analyses were performed separately in MDS and AML, no individual feature remained significantly associated with multi-hit *STAG2* status, likely reflecting the limited size of these subgroup cohorts. Together, these findings support increased dysplasia as the dominant phenotypic correlate of recurrent *STAG2* acquisition while suggesting that the broader clinical phenotype is not driven by differences in baseline demographic or hematologic characteristics.

**Figure 3.**
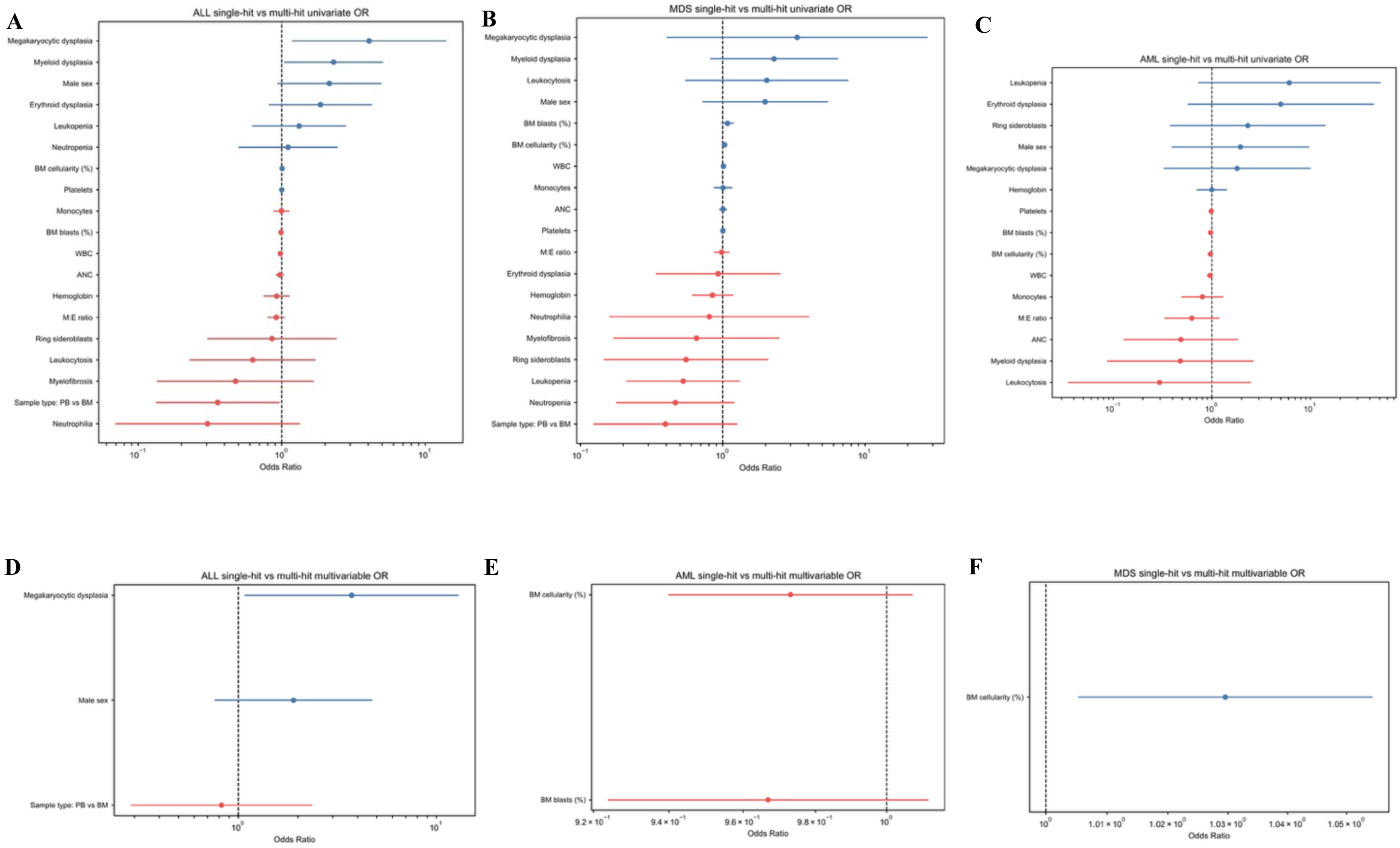
Univariable and multivariable odds of clinical features associated with single versus multiple hits. Forest plot demonstrating the univariable and multivariable odds of the association of various clinical parameters with multiple-hit STAG2 versus single-hit STAG2 in all cases - **A)** & **D)**, MDS cases only - **B)** & **E)**, and AML cases only - **C)** & **F),** respectively.

### Recurrent acquisition of STAG2 mutations occurs throughout clonal evolution

Forty percent of the first-hit mutations in multiSTAG2^MT^ cases were ancestral (dominant: 24%; co-dominant: 16%), while 60% were secondary to other somatic mutations (Figure 4A). The median VAF for STAG2^DOM^ was 63.2% versus 21.1% in STAG2^COD^ and 32.6% in STAG2^SEC^ (Figure 4B). Among the multiSTAG2^MT^ cohort, when the first *STAG2* hit was ancestral (dominant/codominant), the most common secondary mutation was a second *STAG2* hit, followed by *ASXL1*, *TET2*, *BCOR*, *DNMT3A*, and *SRSF2* (Figure 4C). When the first *STAG2* hit was subclonal, the preceding ancestral clone was characterized by *SRSF2*, *ASXL1*, *EZH2*, *BCOR,* and *RUNX1*. Further examination of co-mutational patterns of non-*STAG2* mutations among multi-STAG2MT showed that *ASXL1*, *SRSF2*, *RUNX1,* and *TET2* were the most common, but the distribution of these mutations did not differ based on the clonal hierarchy of the first *STAG2* hit (Figure 4D; eFigure 1). Furthermore, the co-mutational burden was similar regardless of clonal hierarchy status (p = 0.20) (Figure 4E). These differences did not remain significant after correction for multiple testing and should be interpreted as exploratory.

**Figure 4.**
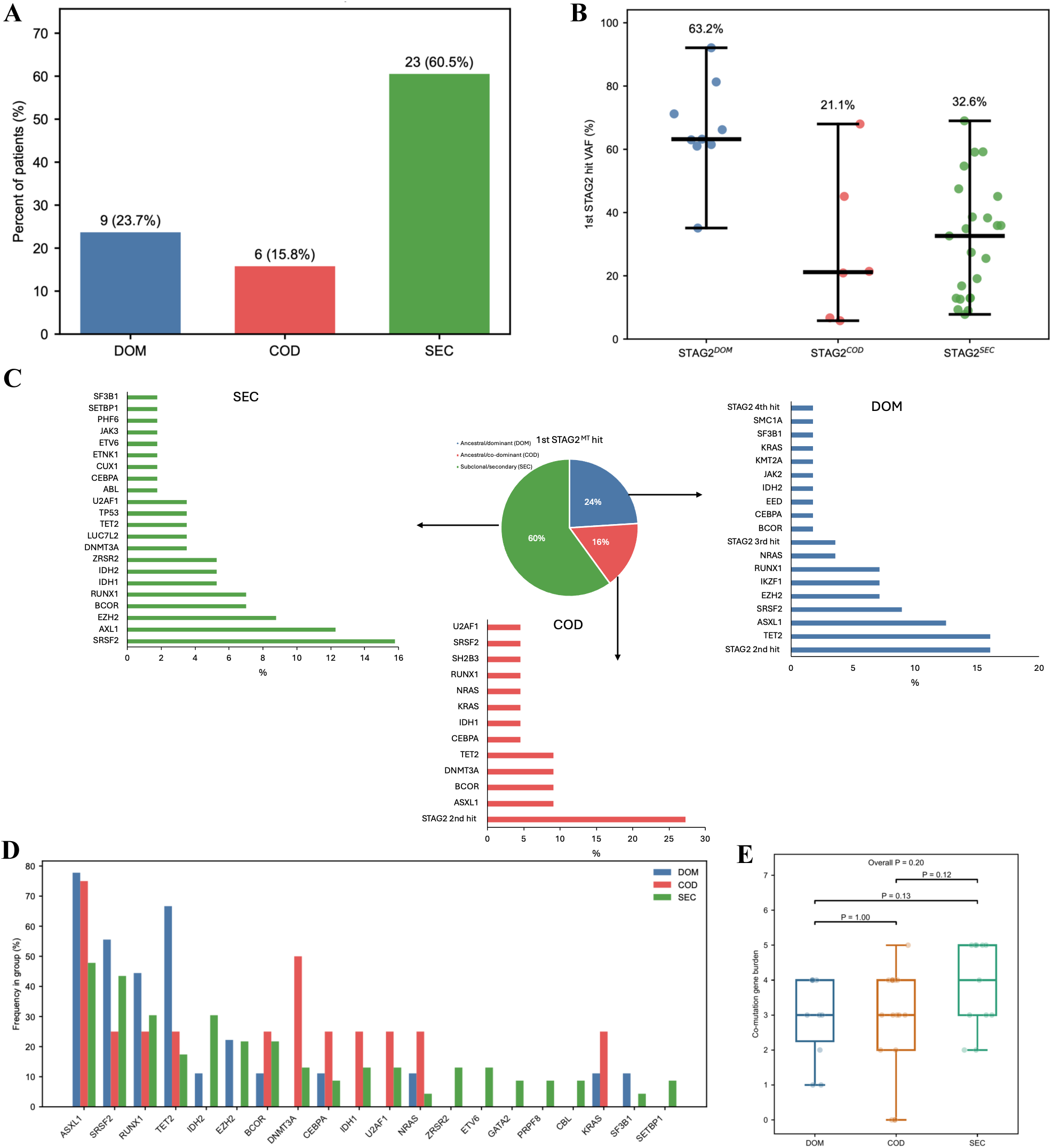
Analysis of STAG2^MT^ features by clonal hierarchy of the 1^st^ STAG2-hit among multiSTAG2^MT^ patients. **A)** Bar graph showing the frequency of clonal hierarchy classification within multiSTAG2^MT^. **B)** VAF distribution of the 1^st^ hit of each multiSTAG2^MT^ patient. **C)** Co-mutational pattern of subsequent or preceding hits for STAG2^MT^ within each clonal ranking. **D)** Bar graph of the co-mutational patterns within each clonal ranking with the exclusion of additional STAG2 hits. **E)** Box plot representing the cumulative mutational gene burden for the 1^st^ STAG2 hit for each clonal rank.

We next examined co-mutational patterns in patients with multiple versus single *STAG2* mutations stratified by disease subtype. In MDS, the largest differences were observed for *IDH1* and *EZH2*, whereas in AML the greatest differences involved *SRSF2* and *RUNX1* (Figure 5A-B). However, none of these associations remained significant after correction for multiple testing, indicating that multi-hit *STAG2* status is not defined by a recurrent cooperating mutational partner. Sex-stratified analyses suggested potential differences in enrichment patterns, including greater representation of *EZH2* mutations among males and *IDH1* mutations among females with MDS, but these findings should be considered exploratory (Figure 5C-D). Collectively, these data suggest that the adverse clinical phenotype associated with multi-hit *STAG2* mutations is not explained by a single recurrent co-mutation.

**Figure 5.**
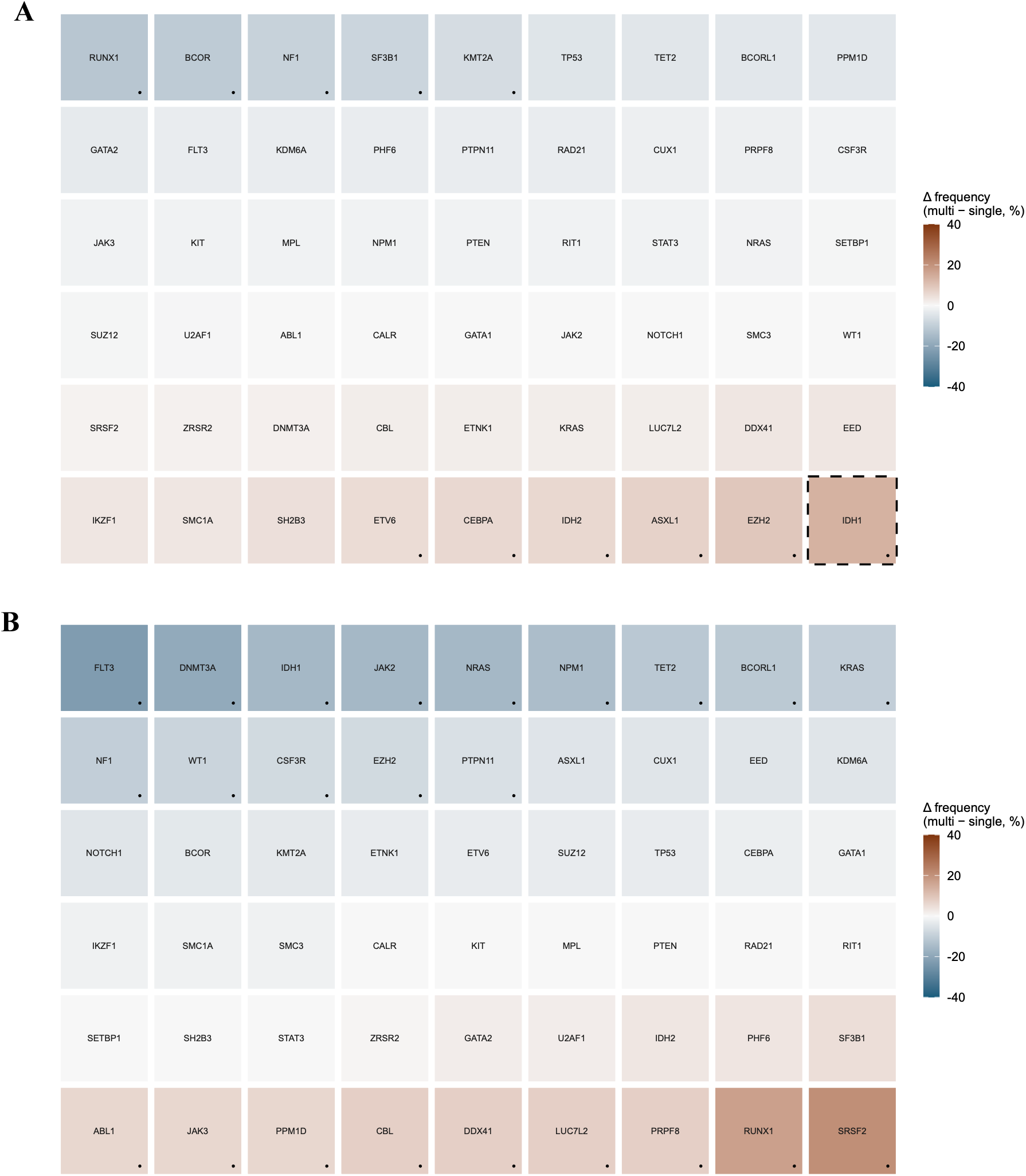
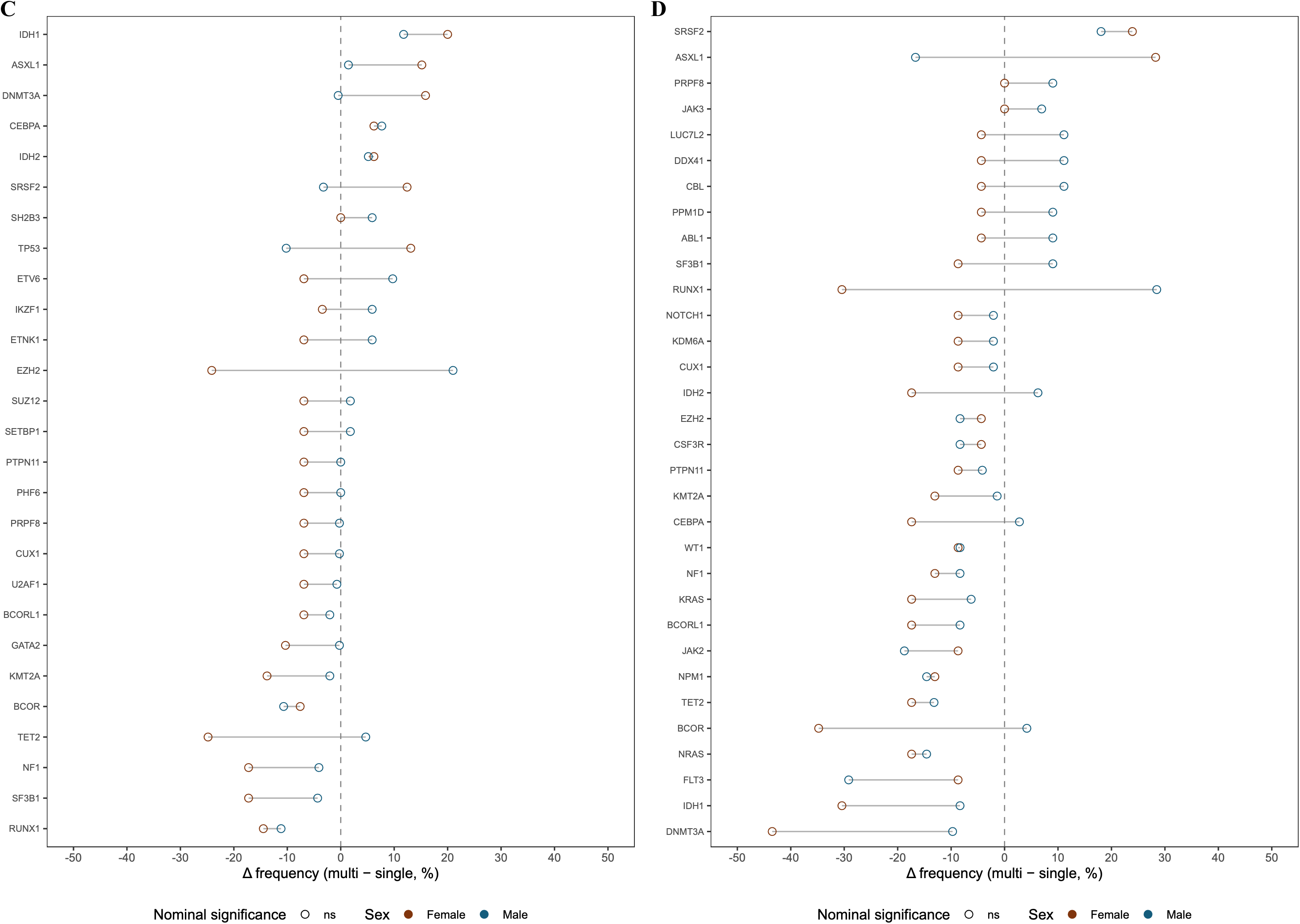
Differences in co-mutational patterns between multiSTAG2^MT^ and monoSTAG2^MT^ patients with MDS and AML. Tile plot showing the difference (Δ) in the frequency of co-mutations (multi-hit minus single-hit *STAG2*) across 54 genes in **A)** MDS patients, and **B)** AML patients. Red tiles indicate enrichment in multi-hit *STAG2* patients while blue tiles indicate enrichment in single-hit *STAG2* patients. Black dots represent cases where |Δ| frequency > 5%. Solid borders represent FDR-adjusted p-values < 0.05; dashed borders represent unadjusted p-values < 0.05. Dumbbell plot demonstrating the difference (Δ) in the frequency of co-mutations in single versus multiple hits per gene in male (blue) and female (red) patients with **C)** MDS and **D)** AML. Each dot represents the difference (relative to zero) in frequency for one sex in a specific gene. The connecting line between the dots represents divergence in enrichment for a specific gene between male and female sex; longer lines represent greater divergence between sexes in the Δfrequency (multi–single). Open circles indicate non-significant differences. All cases indicate instances where |Δ| frequency is > 5% in either sex.

We subsequently stratified multiSTAG2^MT^ cases by their disease subtype to assess differences in the demographic and clinical features based on the clonal hierarchy of the first STAG2 hit. MultiSTAG2^MT^ patients with MDS whose first *STAG2* hit was DOM had a higher WBC and ANC compared to those with COD and SEC clonal ranks (p = 0.01) and resulting leukocytosis (p = 0.04) (eFigure 2). On the contrary, COD multiSTAG2^MT^ patients with MDS had a higher frequency of leukopenia (p = 0.01) and neutropenia (p = 0.01) compared to those with a DOM status. In contrast, overall findings for multiSTAG2^MT^ patients with AML based on clonal rankings were unremarkable (eFigure 3).

### Long-read sequencing supports convergent acquisition of independent STAG2 mutations

Although the majority of multihit cases contained truncating mutations, the temporal ordering and functional interactions between individual STAG2 lesions cannot be resolved from the present dataset. However, the allelic configuration of recurrent STAG2 mutations—specifically whether mutations reside on the same DNA molecule (cis) or arise independently on different alleles or in distinct clones—can be directly assessed using long-read sequencing. To determine the molecular configuration representative of multi-hit *STAG2* events more broadly, we evaluated the feasibility of long-read phasing across additional cases. Among available samples, only two patients (UPN32 and UPN33) harbored mutations separated by genomic distances amenable to single-molecule phasing using long-read sequencing, given the effective read length constraints (∼10–30 kb) and the requirement that all mutations reside within the span of an individual read. In UPN32, two truncating *STAG2* mutations were located within phaseable proximity and demonstrated co-occurrence on different sequencing reads, demonstrating that the mutations did not occur in cis (Figure 6A). In contrast, although UPN33 harbored two *STAG2* mutations, insufficient numbers of single molecules spanning both loci simultaneously (despite adequate locus-level coverage) precluded definitive read-level phasing (Figure 6B).

**Figure 6.**
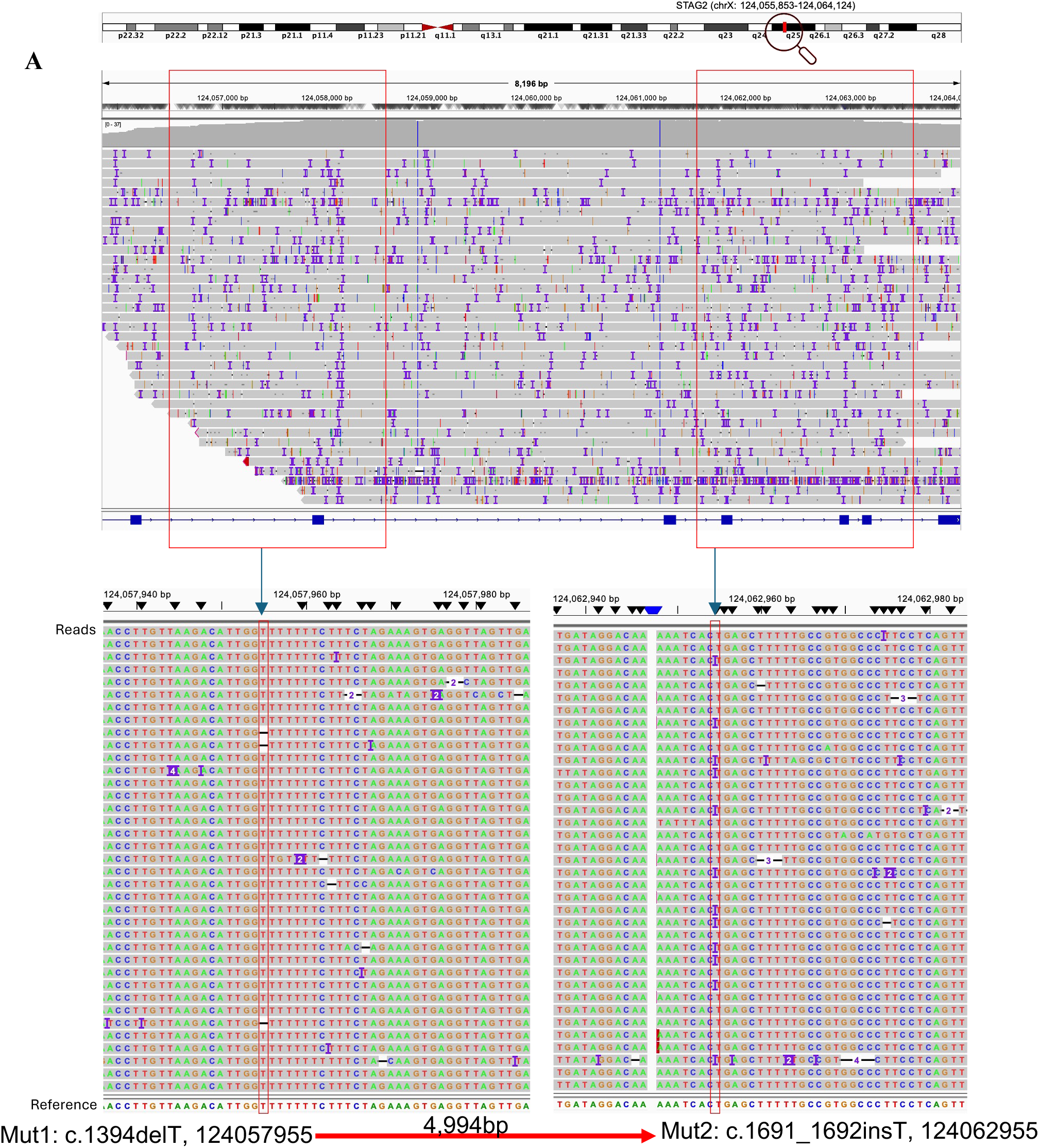
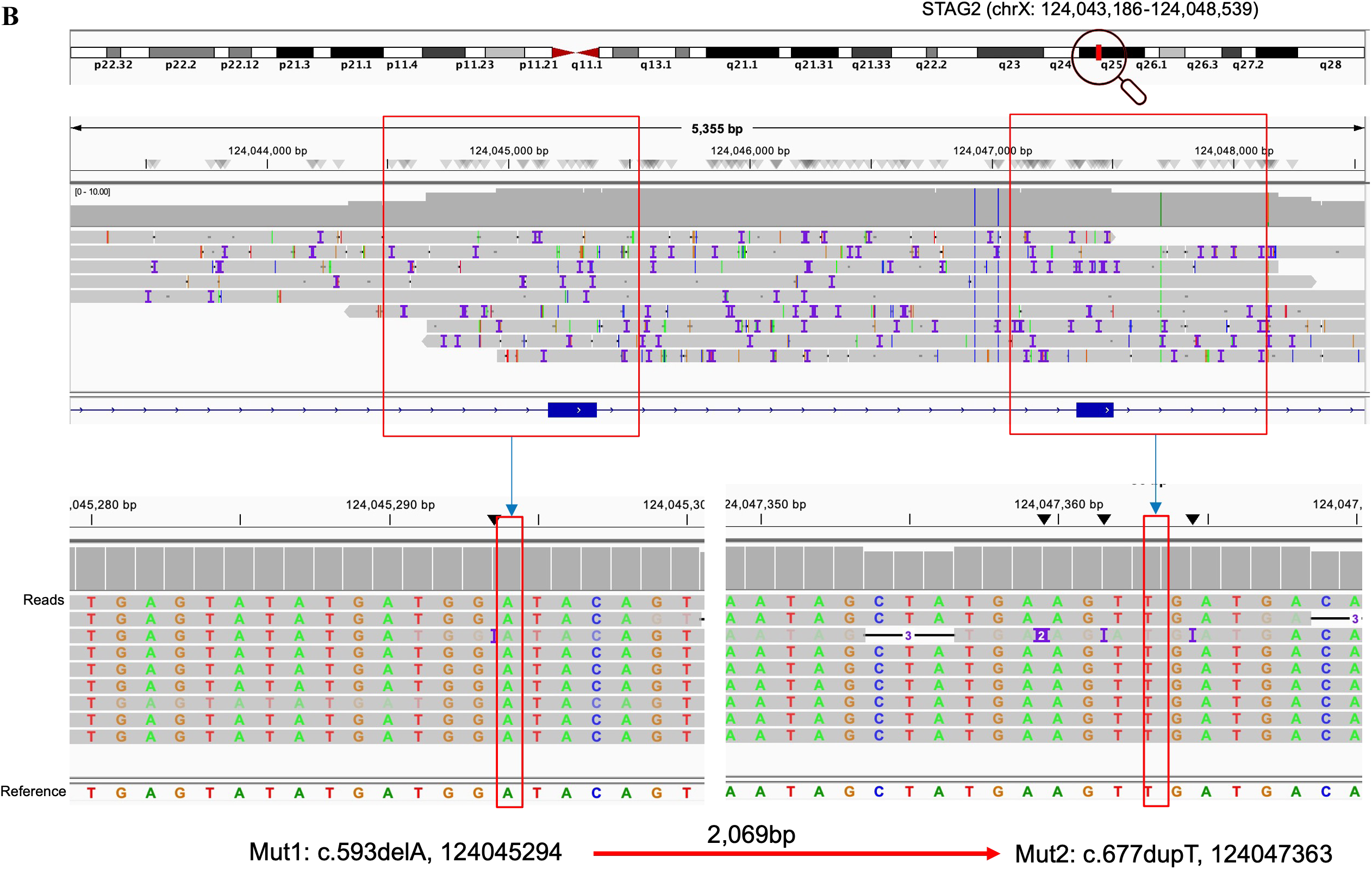
Integrative Genomics Viewer (IGV) output of alignment of long-read nanopore sequencing data from two multiSTAG2^MT^ patient samples. A) UPN 32, and B) UPN 33. Reference genome: GRCH38 – hg 38

By clinical NGS, UPN 32 was reported to harbor three mutations at the indicated genomic position and VAF, respectively: Mut 1 - c.1394delT, 124057961, 12.8%, and Mut 2 - c.1691_1692insT, 38.6%, 124062955, and Mut 3 - c.3133C>T, 4.8%, 124086626. However, only the genomic distance between Mut 1 and Mut 2 was located within phaseable proximity (∼5 kb apart). The presence of these mutations was validated by review of the original DNA tracings from short-read NGS by an independent pathologist (eFigure 4A). Analysis of long-read nanopore sequencing showed that 36 reads overlapped both genomic regions of interest. At genomic position 1, there were no deletions of T observed. However, examining all reads (i.e., not just overlapping reads) revealed the adjusted genomic position to be 124057955, where there were 3 deletions of T (3/36; VAF = 8.3%) (Figure 6A). At genomic position 2 (Mut 2), there were 12 insertions of T (12/36; VAF = 33.3%) (Figure 6A). Deep sequencing analysis demonstrated that multiple STAG2 mutations do not co-occur within the same allele, supporting an independent clonal origin. Instead, these mutations exhibit patterns consistent with independent acquisition, supporting a model of convergent evolution rather than stepwise allelic inactivation.

To complement these findings, we examined variant allele frequency (VAF) relationships among *STAG2* mutations across the entire multi-hit cohort (eTable 4). In most cases, multiple *STAG2* variants demonstrated closely clustered VAFs without clear evidence of mutually exclusive subclonal structures, consistent with these mutations residing within the same dominant clone. Similar VAFs among distinct STAG2 mutations do not necessarily imply a shared allelic origin and may instead reflect expansion of related subclones subjected to similar selective pressures. Accordingly, VAF relationships alone cannot distinguish cis evolution from convergent subclonal acquisition. The direct sequencing data therefore provide the strongest evidence regarding mutational configuration. While some cases demonstrated similar variant allele frequencies across STAG2 mutations, this pattern can be equally explained by independent mutations arising in closely related subclones under strong selective pressure, rather than requiring a cis configuration.

### MultiSTAG2^MT^ status is associated with inferior survival specifically in patients with MDS

Overall, survival is poorer among patients with multiple STAG2 hits compared to patients with a single STAG2 hit (p = 0.04) (Figure 7A). When stratified by disease subtype, this difference is not observed in AML patients (p = 0.83) (Figure 7C). However, MDS patients with multiple hits had poorer outcomes than those with single-hits (p = 0.002) (Figure 7B). This difference in survival persisted regardless of whether the first hit was DOM, COD, or SEC (p = 0.02) (eFigure 5B). Notably, this association was restricted to patients with MDS, with no survival difference observed in AML.

**Figure 7.**
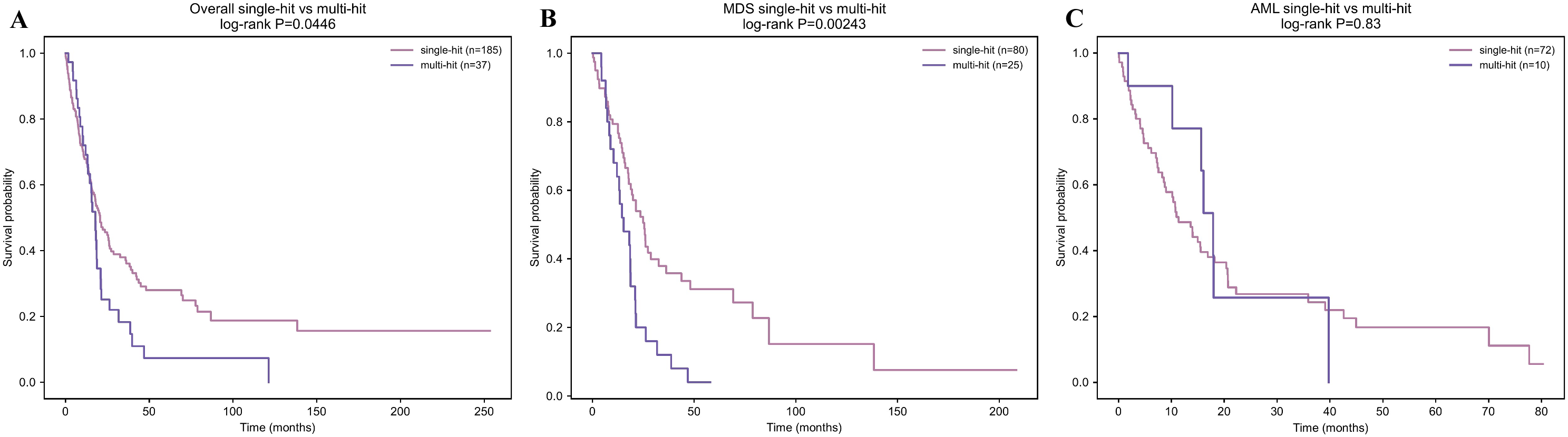
Overall survival analysis in STAG2^MT^ cases by disease subtype. Kaplan-Meier curves comparing overall survival in **A)** all MN cases, **B)** MDS cases only, and **C)** AML cases only.

## Discussion

In this study, we define a distinct subgroup of *STAG2*-mutant myeloid neoplasms characterized by multiple mutations within the same gene and associated with adverse clinical outcomes in MDS. Patients with multi-hit *STAG2* mutations exhibited increased multilineage dysplasia compared with single-hit cases, supporting a model in which cumulative disruption of cohesin function may contribute to disease severity. Our findings support a model in which *STAG2* is repeatedly targeted by independent mutational events across distinct clones, reflecting convergent evolutionary pressure to achieve cohesin dysfunction.

While *STAG2* is classically considered a monoallelic tumor suppressor due to its X-linked location and the predominance of loss-of-function/truncating mutations^24,25^, our data suggest that additional hits are neither incidental nor biologically neutral. Multi-hit states in genes recurrently mutated in MN, including *CEBPA*, *TP53*, and *TET2,* have been associated with distinct clinical phenotypes and variably prognostic impact^13,18,26^. In this context, the adverse outcomes observed in MDS patients with multiple STAG2 mutations are consistent with a gene-dosage model, in which incremental disruption of cohesin function contributes to disease progression beyond a single mutational event^11,12,27^. The absence of cis configuration fundamentally alters the interpretation of multihit *STAG2* states. Rather than reflecting progressive allelic inactivation, these findings indicate that *STAG2* is repeatedly targeted through independent mutational events. This pattern is most consistent with convergent evolution, in which distinct subclonal populations independently acquire mutations in the same gene under shared selective pressure.

Although the prognostic impact of monoallelic *STAG2* mutation is well established, less is known about its clonal architecture in the setting of multiple hits^11^. Using a VAF-based bioanalytic approach, we found that the initial *STAG2* mutation most frequently arose as a secondary rather than ancestral event. This is consistent with prior observations that cohesin mutations occur at multiple stages of leukemogenesis, both as early MDS-defining and progression-associated alleles following early clonal hematopoiesis mutations such as *ASXL1* or *TET2* ^35,36^. The high frequency of a second hit in the ancestral clones may reflect increased permissiveness to subsequent mutational acquisition following initial cohesin perturbation^30^. Notably, the adverse prognostic effect of multi-hit *STAG2* mutations was independent of clonal hierarchy, suggesting that the cumulative allelic burden of *STAG2* disruption—rather than the timing of the initial event—is a key determinant of disease phenotype.

To further resolve the clonal basis of multihit *STAG2* mutations, we applied long-read sequencing to assess allelic configuration. In an informative patient, independent *STAG2* mutations were not observed on the same DNA molecule despite adequate sequencing depth, arguing against a cis configuration. Instead, the data support acquisition of *STAG2* mutations in separate clones, indicating convergent evolutionary targeting of the same gene. These findings suggest that repeated perturbation of *STAG2* provides a selective advantage during disease evolution, even when individual mutations arise independently^30^. Rather than reflecting serial mutational accumulation within a single lineage, this pattern indicates repeated selection for *STAG2* dysfunction across distinct evolutionary trajectories^31–33^.

The biological interpretation of multiple *STAG2* mutations may differ between male and female patients because *STAG2* resides on the X chromosome. In males, recurrent *STAG2* mutations arise within a hemizygous genomic context and therefore reflect repeated targeting of a single functional locus. In females, interpretation is potentially complicated by X-chromosome inactivation and allele-specific clonal selection. Although multihit *STAG2* cases were enriched among males in our cohort, the present study was not designed to evaluate patterns of X-chromosome inactivation, and therefore cannot distinguish between complete or partial lyonization as contributors to STAG2 mutational evolution.

Importantly, prior work has demonstrated that cohesin function is highly dose-dependent. Complete loss of cohesin complex activity is not tolerated in hematopoietic cells, with biallelic deletion resulting in loss of stem cell function and bone marrow failure, whereas partial loss enhances self-renewal and leukemogenesis^34^. This principle is further supported by studies of cohesin paralogs, in which loss of either *STAG1* or *STAG2* alone is compatible with hematopoietic viability, but combined loss results in rapid lethality and marrow aplasia^35,36^. Taken together, these observations support a model in which *STAG2* mutations operate within a constrained, dose-dependent window, whereby incremental loss of function promotes clonal fitness, but complete inactivation is not viable. This framework is consistent with emerging data supporting domain-specific effects of cohesin perturbation and supports a continuum model of tumor suppressor inactivation, ranging from hypomorphic to complete loss-of-function states. These constraints may contribute to the recurrent selection of multi-hit *STAG2* states observed in human disease.

Although direct phasing was feasible in only a limited number of cases, the available evidence supports independent acquisition of *STAG2* mutations rather than compound allelic disruption. Larger studies incorporating routine long-read sequencing will be required to determine how frequently convergent *STAG2* evolution occurs in myeloid neoplasms. Clinically, our findings identify multi-hit *STAG2* mutations as a subgroup of patients enriched for adverse-risk clinical and pathologic features and inferior outcomes in MDS. The association with increased dysplasia and inferior survival suggests that the mutational complexity within *STAG2* may have value for risk stratification and may inform future therapeutic interventions.

The recurrent acquisition of multiple independent *STAG2* mutations suggests that cohesin dysfunction is under strong and sustained selective pressure during disease evolution. Unlike classical tumor suppressors in which biallelic inactivation represents a terminal state, *STAG2* appears to occupy a constrained functional window in which partial impairment is advantageous, but complete loss is disfavored. Convergent targeting of *STAG2* across subclones therefore represents a mechanism for achieving this optimal level of dysfunction.

In summary, multi-hit *STAG2* mutations define a biologically and clinically distinct subset of myeloid neoplasms characterized by convergent evolutionary targeting of *STAG2*. These findings highlight the importance of clonal architecture in interpreting mutations in X-linked tumor suppressors and suggest that repeated selection for cohesin dysfunction contributes to adverse disease biology.

## Acknowledgments

1. E. B. U. is supported by the American Society of Hematology (ASH) Physician Scientist (Phy- Sci) Award. A.D.V is supported by an ASH Scholar award and by the Gabrielle’s Angels Foundation. A.D.V. is a Damon-Runyon-Doris Duke Clinical Investigator supported by the Damon Runyon Cancer Research Foundation (CI-120-22) and a Clinician Scientist Development Award from the Doris Duke Charitable Foundation. This research was funded in part through the NIH/NCI Cancer Center Support Grant (P30CA012696), supported by the NCATS Clinical and Translational Science Awards (CTSA) grant (UL1TR001873), and used the Genomics and High Throughput Screening Shared Resource.

Schematics included in the figures were made using BioRender.

## Authorship contributions

1. E. B. U. – conceptualization/design, data curation, investigation, analysis, visualization, writing – original draft, writing – review and editing. Y.C – data curation, visualization, analysis. M.D. - conceptualization/design, data curation, investigation. E.S. & R.D. – technical/methodological support, analysis. S.U. & Z.B. – data curation. V.S. – data visualization. K.T. & D.S. – technical support, data curation. V.V. & J.P.M. – conceptualization/design, investigation, methodology. A.D.V. – conceptualization/design, investigation, funding acquisition, methodology, project supervision, writing – original draft, writing – review and editing. All other authors reviewed the clinical data and helped in editing the manuscript. All authors provided data interpretation and review of the final manuscript.

