## Supplementary Materials for "Multi-hit STAG2 mutations define a high-risk subset of MDS and reveal convergent evolutionary targeting of cohesin"

**Supplementary Material**

**eTable 1. List of genes covered by Cleveland Clinic hematologic neoplasm next-generation sequencing (NGS) panel.**

| | | | | | | | |
|---|---|---|---|---|---|---|---|
| ABL1 | ASXL1 | BCOR | BCORL1 | BRAF | CALR | CBL | CDKN2A |
| CEBPA | CSF3R | CUX1 | DDX41 | DNMT3A | EED | ETNK1 | ETV6 |
| EZH2 | FBXW7 | FLT3 | GATA1 | GATA2 | GNAS | IDH1 | IDH2 |
| IKZF1 | JAK2 | JAK3 | KDM6A | KIT | KMT2A | KRAS | LUC7L2 |
| MPL | MYD88 | NF1 | NOTCH1 | NPM1 | NRAS | PAX5 | PHF6 |
| PIGA | PPM1D | PRPF8 | PTEN | PTPN11 | RAD21 | RIT1 | RUNX1 |
| SETBP1 | SF3B1 | SH2B3 | SMC1A | SMC3 | SRSF2 | STAG2 | STAT3 |
| STAT5B | SUZ12 | TET2 | TP53 | U2AF1 | WT1 | ZRSR2 | |

**eTable 2. Comparison of baseline characteristics in multiSTAG2$^{MT}$ and monoSTAG2$^{MT}$ MN cases.**

| Variables | monoSTAG2$^{MT}$ (n = 195) | multiSTAG2$^{MT}$ (n = 38) | p-value |
|---|---|---|---|
| **Age (y)** | 73.7 (65.5, 80.7) | 71.3 (65.6, 78.8) | 0.67 |
| ≥60 | 159 (81.5) | 34 (89.5) | 0.35 |
| <60 | 36 (18.5) | 4 (10.5) | |
| **Gender** | | | 0.07 |
| Male | 124 (63.6) | 30 (79.0) | |
| Female | 71 (36.4) | 8 (21.0) | |
| **WHO Classification (2016)** | | | |
| **MDS** | 78 (40.0) | 22 (57.9) | 0.30 |
| MDS-EB-1 | 25 | 5 | |
| MDS-EB-2 | 14 | 10 | |
| MDS-MLD-RS | 4 | 3 | |
| MDS-U | 2 | 1 | |
| MDS-MLD | 28 | 3 | |
| **MDS/MPN** | 12 (6.2) | 3 (7.9) | |
| CMML | 9 | 1 | |
| MDS/MPN-U | 1 | 2 | |
| **AML** | 71 (36.4) | 11 (29.0) | |
| sAML | 35 | 8 | |
| tAML | 0 | 1 | |
| pAML | 34 | 2 | |
| **MPN** | 7 (3.6) | 0 (0.0) | |
| MF | 2 | 0 | |
| ET/MF | 1 | 0 | |
| PV | 1 | 0 | |
| **Other** | 27 (13.9) | 2 (5.3) | |
| CH | 5 | 0 | |
| CCUS | 20 | 2 | |
| **Cytogenetics** | | | 0.50 |
| Normal karyotype | 100 (62.1) | 19 (55.9) | |
| Abnormal karyotype | 61 (37.9) | 15 (44.1) | |

**eTable 3. Comparison of clinical characteristics in multiSTAG2$^{MT}$ and monoSTAG2$^{MT}$ MN cases.**

| Variables | monoSTAG2$^{MT}$ (n = 195) | multiSTAG2$^{MT}$ (n = 38) | p-value |
|---|---|---|---|
| **Hematologic parameters** | | | |
| WBC ($10^9$/L) (median/range) | 2.0 (1.0, 5.0) | 1.0 (1.0, 4.0) | 0.42 |
| <4.5 x $10^9$/L - leukopenia | 112 (60.22) | 24 (66.7) | 0.47 |
| >11.0 x $10^9$/L - leukocytosis | 38 (20.4) | 5 (13.9) | 0.36 |
| Hemoglobin (g/dL) | 8.6 (7.6, 9.7) | 8.0 (7.5, 9.5) | 0.27 |
| <12 g/dL (F) or <13g/dL (M) - anemia | 179 (94.2) | 35 (97.2) | 0.70 |
| >17 g/dL - erythrocytosis | 1 (0.54) | 0 (0.0) | 1.00 |
| Platelet ($10^9$/L) | 68 (36, 139) | 64 (29, 120) | 0.48 |
| <150 X $10^9$/L - thrombocytopenia | 144 (77.4) | 30 (83.3) | 0.43 |
| >450 X $10^9$/L - thrombocytosis | 9 (4.8) | 2 (5.6) | 0.69 |
| ANC ($10^9$/L) | 1.39 (0.43, 4.28) | 0.85 (0.34, 2.67) | 0.12 |
| <1.8 x $10^9$/L - neutropenia | 115 (61.8) | 20 (64.5) | 0.87 |
| >6.0 x $10^9$/L - neutrophilia | 30 (16.1) | 2 (5.6) | 0.26 |
| **Cytopenias** | | | |
| Monocytopenia | 25 (13.6) | 4 (11.1) | 0.66 |
| Bicytopenia | 92 (50.0) | 11 (30.6) | |
| Pancytopenia | 67 (36.4) | 21 (58.3) | |
| **Bone Marrow Morphology** | | | |
| Blasts | 3 (3, 30) | 9 (3, 18) | 0.52 |
| >= 5% | 104 (53.3) | 23 (60.5) | 0.42 |
| Cellularity | | | 0.45 |
| Hypercellular | 91 (58.0) | 19 (55.9) | |
| Normocellular | 37 (23.6) | 11 (32.4) | |
| Hypocellular | 29 (18.5) | 4 (11.8) | |
| M/E ratio | 1.7 (1.1, 3.8) | 2.6 (1.2, 4.8) | 0.28 |
| Myelofibrosis | 29 (18.4) | 3 (9.7) | 0.30 |
| **Dysplastic Lineage** | | | |
| Myeloid | 74 (45.4) | 21 (65.6) | **0.04** |
| Erythroid | 96 (58.9) | 24 (72.7) | 0.14 |
| Megakaryocytic | 112 (70.4) | 29 (90.6) | **0.02** |
| **Number of dysplastic BM lineage** | | | |
| None | 30 (18.3) | 0 (0.0) | **0.02** |
| Unilineage | 28 (17.1) | 4 (12.5) | |
| Bilineage | 64 (39.0) | 14 (43.8) | |
| Trilineage | 42 (25.6) | 14 (43.8) | |

**eTable 4. Variant allele frequencies of each *STAG2* mutation within the multi-hit cohort.**

| UPN | VAF 1 | VAF 2 | VAF 3 | VAF 4 |
|---|---|---|---|---|
| 1 | 12.6 | 4.0 | | |
| 2 | 18.7 | 19.1 | | |
| 3 | 35.9 | 9.4 | | |
| 4 | 11.2 | 6.1 | 63.2 | 13.6 |
| 5 | 5.4 | 12.9 | | |
| 6 | 20.9 | 6.7 | | |
| 7 | 4.2 | 38.3 | | |
| 8 | 5.3 | 12.9 | 4.2 | |
| 9 | 4.4 | 6.7 | | |
| 10 | 7.8 | 61.0 | | |
| 11 | 6.7 | 16.8 | | |
| 12 | 92.1 | 5.4 | | |
| 13 | 27.4 | 7.5 | | |
| 14 | 32.6 | 6.4 | 6.5 | 6.8 |
| 15 | 5.2 | 5.8 | | |
| 16 | 13.2 | 81.3 | | |
| 17 | 12.1 | 45.1 | | |
| 18 | 10.7 | 66.2 | | |
| 19 | 71.2 | 13.0 | | |
| 20 | 21.4 | 15.5 | | |
| 21 | 17.9 | 61.5 | | |
| 22 | 9.0 | 63.0 | 14.0 | |
| 23 | 6.0 | 6.0 | 9.0 | |
| 24 | 11.0 | 13.0 | 13.0 | |
| 25 | 40.4 | 47.5 | | |
| 26 | 35.9 | 9.4 | | |
| 27 | 4.3 | 45.1 | | |
| 28 | 35.1 | 5.8 | | |
| 29 | 13.6 | 25.5 | | |
| 30 | 7.6 | 7.8 | | |
| 31 | 5.6 | 69.0 | | |
| 32 | 4.8 | 38.6 | 12.8 | |
| 33 | 31.2 | 59.1 | | |
| 34 | 7.2 | 59.2 | | |
| 35 | 54.7 | 17.9 | | |
| 36 | 4.3 | 68.0 | | |
| 37 | 34.9 | 10.2 | | |
| 38 | 9.3 | 6.0 | | |

UPN – unique patient number, VAF – variant allele frequency.

### Supplementary Figure Legends

**eFigure 1. Clonal hierarchy of the first STAG2 mutation was not associated with reproducible differences in co-mutational patterns, supporting recurrent STAG2 acquisition across diverse genetic backgrounds.** Forest plot showing the odds ratios (OR) of genes commonly mutated in myeloid neoplasms among multiSTAG2$^{MT}$ cases for **A)** DOM vs. SEC. **B)** COD vs SEC. **C)** DOM vs. COD clonal rankings.

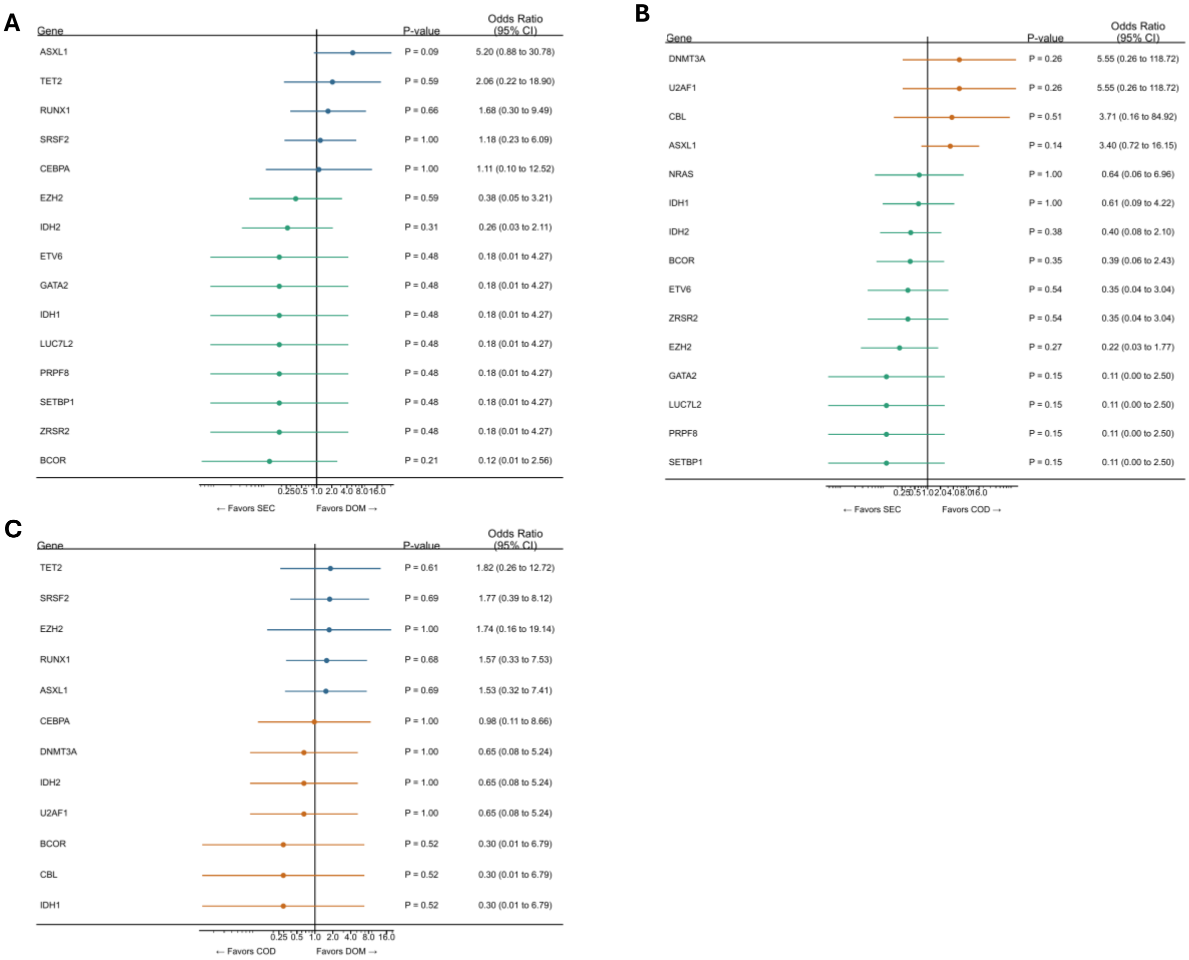

**eFigure 2. Among MDS patients with multi-hit STAG2 mutations, dominant first-hit STAG2 clones were associated with higher leukocyte counts, whereas codominant clones were enriched for leukopenia and neutropenia.** Box plots displaying the levels of **A)** WBC, **B)** Hemoglobin, **C**) Platelet, **D)** ANC, **E)** BM blast %, **F)** BM cellularity, **G)** Monocytes, and **H)** M/E ratio in multiSTAG2$^{MT}$ cases with MDS based on the clonal status of the 1st STAG2 hit.

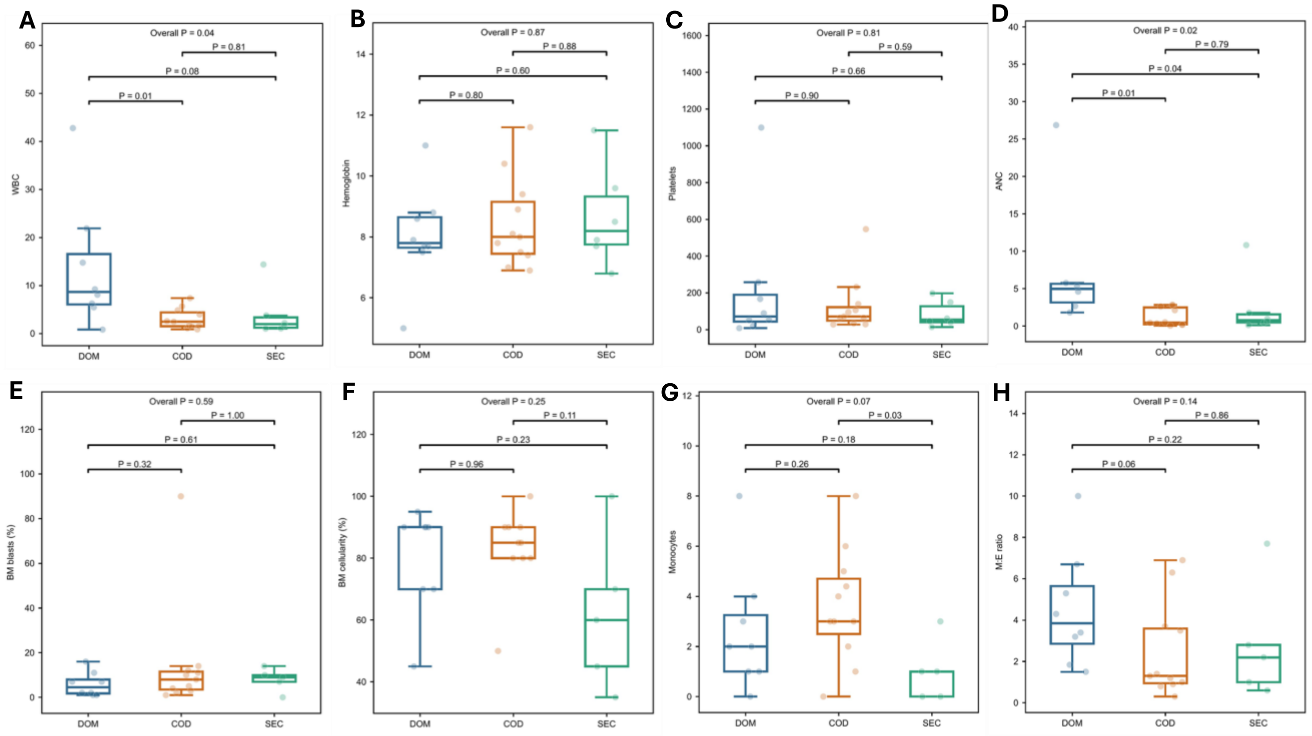

**eFigure 3. Clinical and hematologic characteristics were largely similar across clonal hierarchy groups in AML, indicating limited phenotypic impact of STAG2 clonal timing in this disease context.** Box plots displaying the levels of **A)** WBC, **B)** Hemoglobin, **C**) Platelet, **D)** ANC, **E)** BM blast %, **F)** BM cellularity, **G)** Monocytes, and **H)** M/E ratio of multiSTAG2$^{MT}$ cases with AML based on the clonal status of the 1st STAG2 hit.

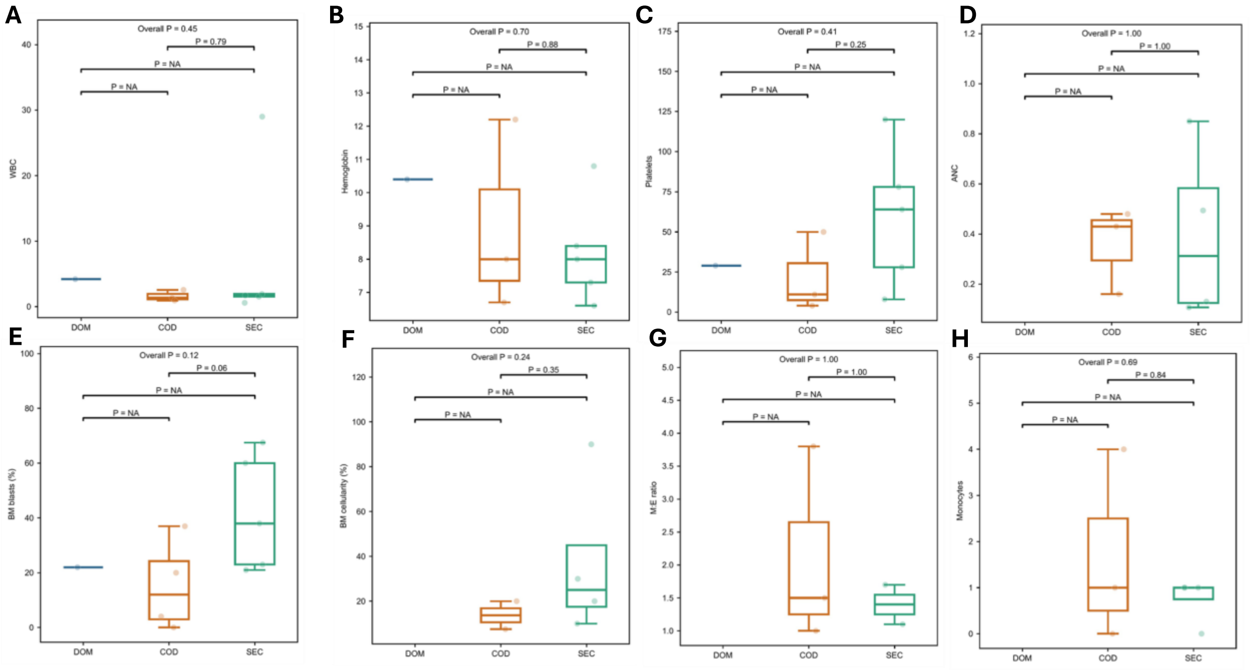

**eFigure 4. Independent review of clinical short-read sequencing data confirmed the presence of multiple STAG2 mutations selected for long-read allelic phasing analyses.** Patient samples aligned to the reference genome (GRCH38 – hg 38)**. A)** UPN 32, and **B)** UPN 33.

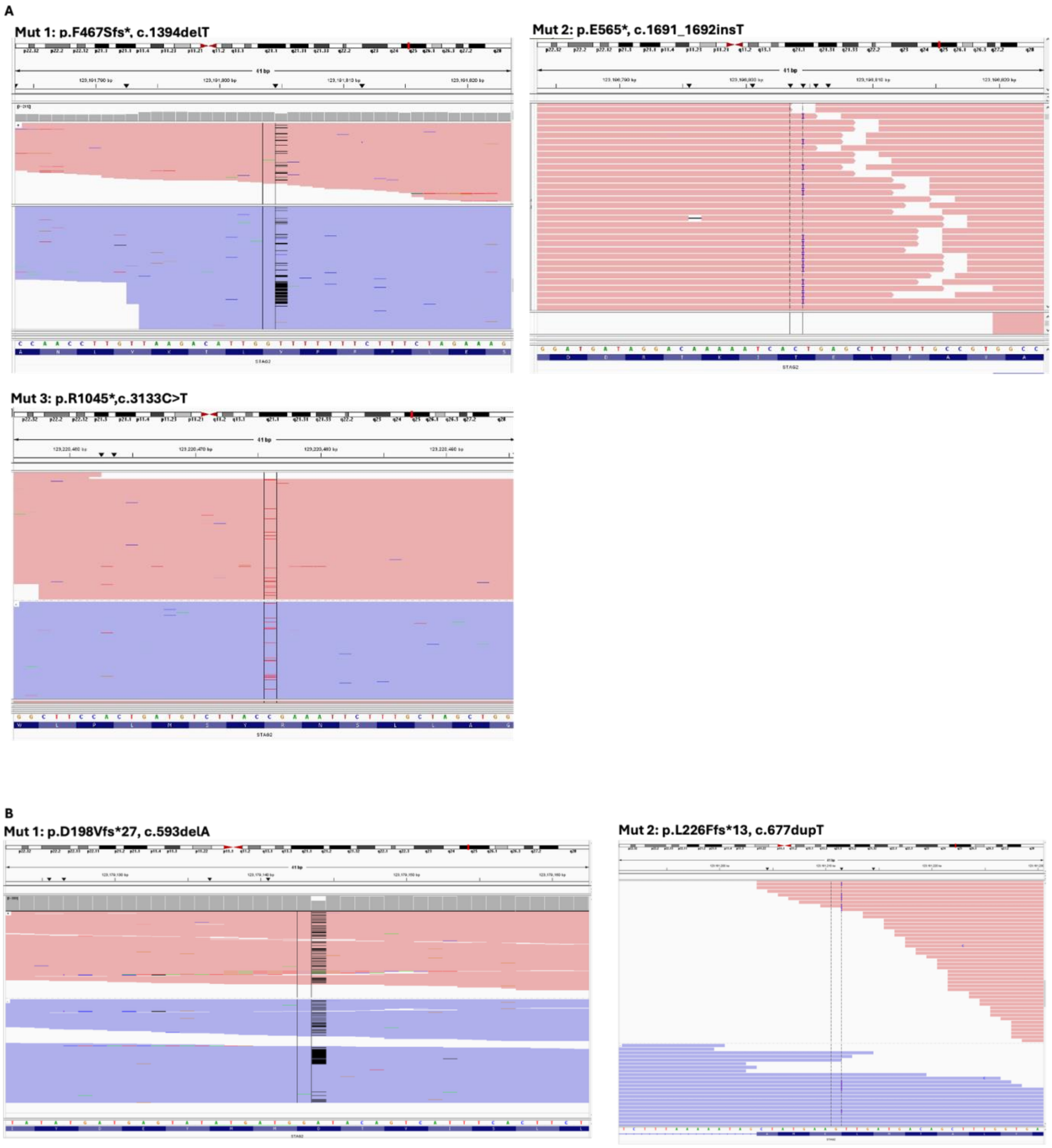

**eFigure 5. The adverse prognostic impact of multi-hit STAG2 mutations was restricted to MDS and was not observed in AML, regardless of clonal hierarchy.** Kaplan-Meier curves comparing overall survival in **A)** all MN cases, **B)** multi-hit STAG2$^{MT}$ MN cases only, **C)** all MDS cases, **D)** multi-hit STAG2$^{MT}$ MDS cases only, **E)** all AML cases, **F)** multi-hit STAG2$^{MT}$ AML cases only.

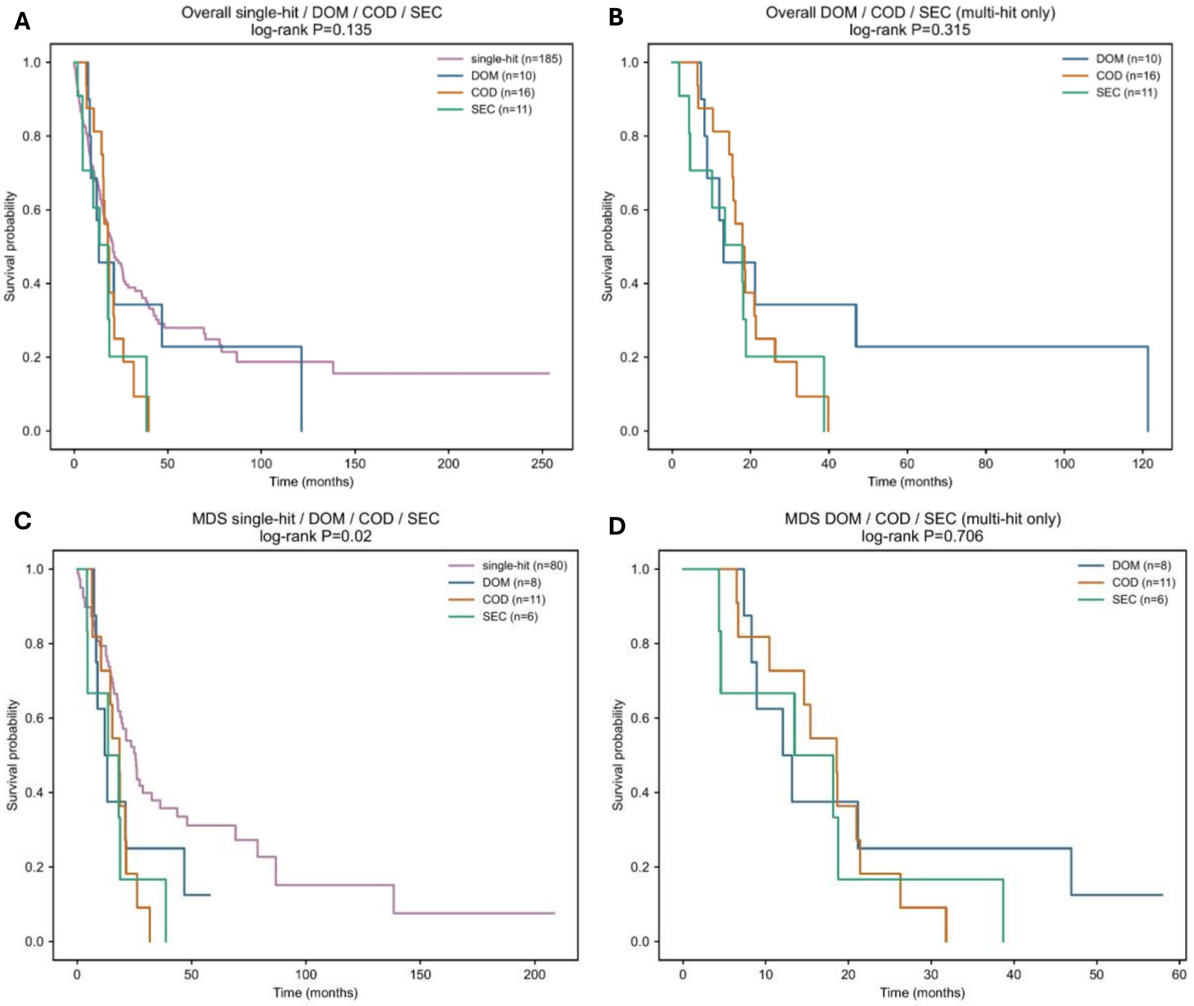

**E**

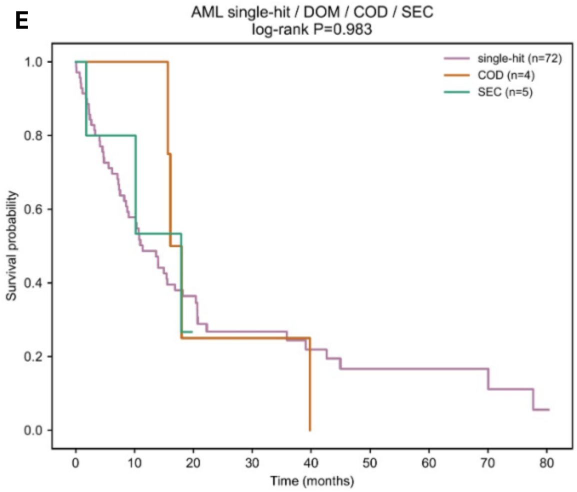
AML single-hit / DOM / COD / SEC
log-rank P=0.983
single-hit (n=72)
COD (n=4)
SEC (n=5)
Survival probability
1.0
0.8
0.6
0.4
0.2
0.0
0
10
20
30
40
50
60
70
80
Time (months)

**F**

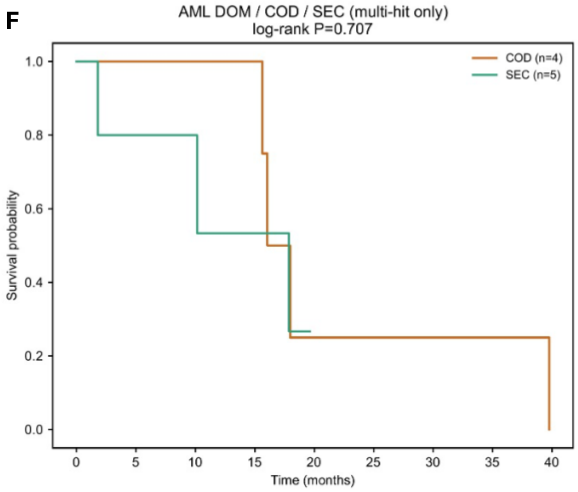
AML DOM / COD / SEC (multi-hit only)
log-rank P=0.707
COD (n=4)
SEC (n=5)
Survival probability
1.0
0.8
0.6
0.4
0.2
0.0
0
5
10
15
20
25
30
35
40
Time (months)
